# Balancing Locality and Reconstruction in Protein Structure Tokenizer

**DOI:** 10.1101/2024.12.02.626366

**Authors:** Jiayou Zhang, Barthelemy Meynard-Piganeau, James Gong, Xingyi Cheng, Yingtao Luo, Hugo Ly, Le Song, Eric Xing

## Abstract

The structure of a protein is crucial to its biological function. With the expansion of available protein structures, such as those in the AlphaFold Protein Structure Database (AFDB), there is an increasing need for efficient methods to index, search, and generate these structures. Additionally, there is a growing interest in integrating structural information with models from other modalities, such as protein sequence language models. We present a novel VQ-VAE-based protein structure tokenizer, AIDO.StructureTokenizer (AIDO.St), which is a pretrained module for protein structures in an AI-driven Digital Organism [1]. AIDO.StructureTokenizer is a 300M parameter model consisting of an equivariant encoder to discretize input structures into tokens, and an invariant decoder to reconstruct the inputs from these tokens. In addition to evaluating structure reconstruction ability, we also compared our model to Foldseek, ProToken, and ESM3 in terms of protein structure retrieval ability. Through our experiments, we discovered an intriguing trade-off between the encoder’s locality and retrieval ability and the decoder’s reconstruction ability. Our results also demonstrate that a better balance between retrieval and reconstruction enables a better alignment between the structure tokens and a protein sequence language model, resulting in better structure prediction accuracy. Models and code are available through ModelGenerator in https://github.com/genbio-ai/AIDO and on Hugging Face.

## 1 Introduction

The rapid developments of protein structure prediction methods such as AlphaFold have resulted in a vast increase in the protein structure databases [2, 3], opening up new avenues for integrating structural information to better understand proteins. Despite these advancements, leveraging the full potential of protein structures within large-scale models remains a significant challenge, particularly due to the complexity and computational demands of working with 3D structural data.

Tokenizing protein structures is an innovative approach designed to convert the complex 3D information of proteins into a discrete, more manageable format that can be seamlessly integrated with sequence-based models. Traditional methods for analyzing protein structures typically rely on processing 3D coordinates or distance matrices, which are computationally intensive and difficult to align with sequence data. Recently, there has been increasing interest in developing models that can incorporate structural information alongside sequence data [4, 5, 6].

In this work, we introduce a novel vector quantization variational autoencoder (VQ-VAE) architecture for protein structure tokenization, featuring an equivariant encoder and an invariant decoder. We trained one of the largest VQ-VAE models to date, utilizing 300 million parameters on over 477,000 protein structures from the PDB. This tokenizer efficiently converts protein structures into discrete tokens, facilitating their seamless integration with sequence-based models.

While most tokenizers focus solely on reconstruction accuracy—which provides only a partial indication of tokenizer quality—we evaluate ours through additional tasks such as homology detection and integration into a protein language model for structure prediction. We observe an interesting trade-off between the encoder locality and the reconstruction ability of the decoder. By comparing to Foldseek, ProToken and ESM3, we showed that our model can achieve a better balance by obtaining substantial improvements in various structure retrieval tasks and prediction tasks while at the same time sacrificing less than 2% in the overall structure reconstruction ability. Our results point to the direction of a more efficient and accurate framework for protein structure modeling.

## 2 Background and Related Work

Foldseek [7] introduced an efficient method for tokenizing protein structures by encoding local features, such as distances and angles, to accelerate homology detection. However, this approach results in significant information loss, limiting its usefulness for tasks that require detailed structural reconstruction [7]. ProToken addressed this limitation by employing a symmetric encoder-decoder architecture, allowing for high-fidelity reconstruction from tokens, though these tokens have proven less effective in broader downstream applications [8]. Subsequent models, such as ProstT5 and SaProt, built on Foldseek’s tokenization to enhance protein language models, improving performance in tasks like mutation effect prediction by integrating structural tokens with sequence data [5]. ProSST further refined this approach by implementing a local denoising autoencoder and expanding the token vocabulary, leading to superior results in mutation effect prediction and underscoring the importance of optimizing tokenization schemes for better downstream performance [6]. ESM3 is one of the most successful applications of VQ-VAE for protein structure tokenization in multimodal modeling [9]. By using VQ-VAE to discretize protein structures into tokens, ESM3 effectively integrates both structural and sequence data within a large language model (LLM). This integration enables conditional sequence generation and structure prediction. Beyond sequence and structure, ESM3 incorporates features and annotations, making it versatile for a wide range of biological tasks.

## 3 Methods

We employ a Vector Quantized Variational Autoencoder (VQ-VAE) to tokenize the 3D structure of proteins. The primary goal of this tokenization is to transform the continuous geometric data of protein backbones into discrete tokens that can be later integrated into sequence-based models.

### 3.1 Architecture Details

The VQ-VAE architecture used in our approach is a novel combination of an equivariant encoder and an invariant decoder (see Fig. 1.A for an overview). More specifically,

**Figure 1.**
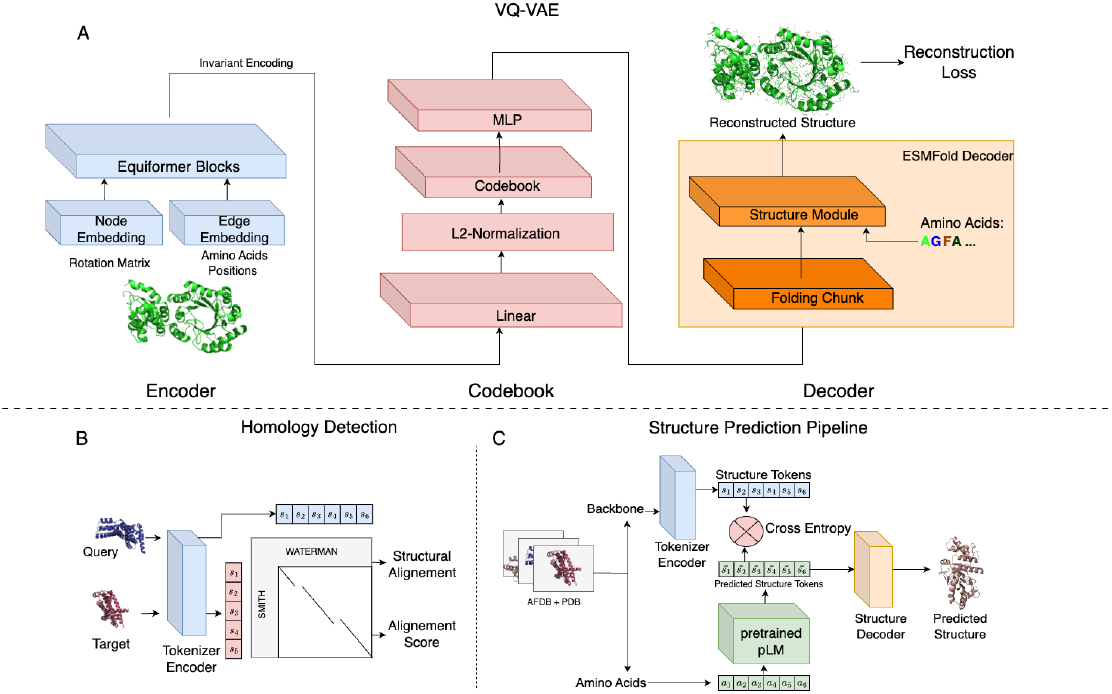
A.The backbone structure is encoded as residue-level frame, comprising a rotation matrix, translation vector, and residue index. The rotation matrix is used as node features, while inter-node translation and residue index differences for edge features. Equiformer blocks propagate information between k-nearest neighbors and outputs processed node embeddings. The invariant part of these embeddings is projected to a lower dimension and discretized via a VQ-VAE Codebook. The Structure Decoder then processes these tokens and reconstructs the full 3D structure, including side chains, using the ESMFold-based Folding Trunk. B. Homology Detection: Structures from the SCOPe40 database are tokenized and aligned using Smith-Waterman. This gives us a score that we can use to evaluate if two proteins are homologous. C. Structure Prediction: We fine-tune a Language Model to predict the structure tokens given by the encoder. The predicted tokens are then given to the decoder which will output the predicted structure.

- **Structure Encoder:** The encoder projects the backbone structure into a latent space where each residue is represented as a vector. The encoder is based on the Equiformer architecture [10], which incorporates local attention mechanisms. To ensure computational efficiency and maintain locality in the extracted features, the attention scope is restricted to the 30-nearest neighbors in the input structure. We choose a rather small encoder (6M) compared to the large decoder (300M) following finding in [11].
- **VQ-VAE Codebook:** The invariant part of the latent vectors produced by the encoder are then quantized by mapping each vector to the nearest entry in a fixed-size codebook following the procedure in [12]. This process converts the continuous latent vectors into discrete tokens, conserving the amino acid sequence length.
- **Structure Decoder:** The quantized vectors are fed into a Multi-Layer Perceptron (MLP), which generates the “single embedding” input for the decoder which reuses the ESMFold Folding Trunk architecture [13]. The Folding Trunk then reconstructs the full 3D structure, including side chains, based on the structural tokens. The reconstruction process is guided by several loss functions, including the Frame Aligned Point Error (FAPE) loss, which measures the discrepancy between the predicted and actual protein structures. All losses and training details are included in Appendix A.3.

We observed that the locality of the encoder—how much local structural information each token captures—significantly impacts both performance and the type of information encoded. Foldseek, for instance, employs a highly local approach by considering only the 2 nearest neighbors (for these two residues it also uses the residues before and after to compute some relative vector, making the actual receptive field between 2 and 6) during encoding. ProToken, on the other hand, uses full attention mechanisms, capturing global dependencies across the entire protein structure. Both ESM and our model strike a balance between these extremes: ESM utilizes a k-nearest neighbors (kNN) value of 16, while our model uses a kNN of 30. In our model kNN means that we mask the attention such that each query can only attend its 30 nearest neighbors. We believe that this choice of locality is a major modeling decision that influences the model’s performance and the nature of the structural information encoded in the tokens.

More details of the architecture and the training methods can be found in the Appendix A.2 and A.3.

### 3.2 Alignment to Protein Language Model

To integrate structural information with a protein language model (pLM), we tokenized the AlphaFold Database (AFDB) using our VQ-VAE-based tokenizer. This produced discrete structural tokens aligned with the amino acid sequences, where the i-th amino acid directly corresponds to the i-th structural token, ensuring a natural alignment between sequence and structure.

We fine-tuned pre-trained pLM to predict these structural tokens from the amino acid sequence using two models: ESM2-650M [13] for efficiency, and our in house MoE pLM [14] for comprehensive model optimization. The predicted tokens can then be passed to the VQ-VAE decoder, transforming the pipeline into a full structure prediction model capable of generating 3D protein structures from sequence-only data (see Fig. 1.C). More details can be found in Appendix A.4.

## 4 Tokenizer Evaluation

Evaluating the effectiveness of a tokenizer is not a straightforward task, as it requires careful consideration of what constitutes a “good” tokenization. In the context of protein structures, a good tokenizer must encapsulate all the essential structural information. However, it is equally important that this information is preserved in a coherent and interpretable manner. Specifically, given the goal of using these tokens for multimodal tasks, such as integrating protein sequences and structures, the tokenization process must maintain the sequentiality of the original data. This means that each token should correspond meaningfully to its respective amino acid in the sequence, ensuring that the *i*-th token remains closely related to the *i*-th amino acid. Maintaining this correspondence is critical for enabling downstream tasks that rely on the alignment of sequence and structure information, such as joint training with protein language models. We evaluate the effectiveness of our approach through a comprehensive set of tasks that assess the quality of the tokenization.

### 4.1 Reconstruction Evaluation

The quality of the tokenization was previously assessed primarily through reconstruction performance, comparing the output of the decoder to the original input structure. This evaluation reflects the effectiveness with which structural information is encoded into discrete tokens as shown in Table 1. The metrics used are TM-score, RMSD (Root Mean Square Deviation), and GDT (Global Distance Test). These metrics provide a quantitative assessment of how closely the reconstructed protein matches the original structure. See the Appendix B.1 for detailed definitions of these metrics. In the reconstruction evaluation, we observed that certain proteins, particularly those with a larger radius (greater than 100 Å), posed a challenge for the model (examples in Appendix C.1). Therefore, we decomposed the performance based on the protein’s radius, separately evaluating proteins with radii below and above 100 Å. This allowed us to analyze how well the tokenizer encodes information across different structural scales and identify areas where further improvements may be needed.

**Table 1:**
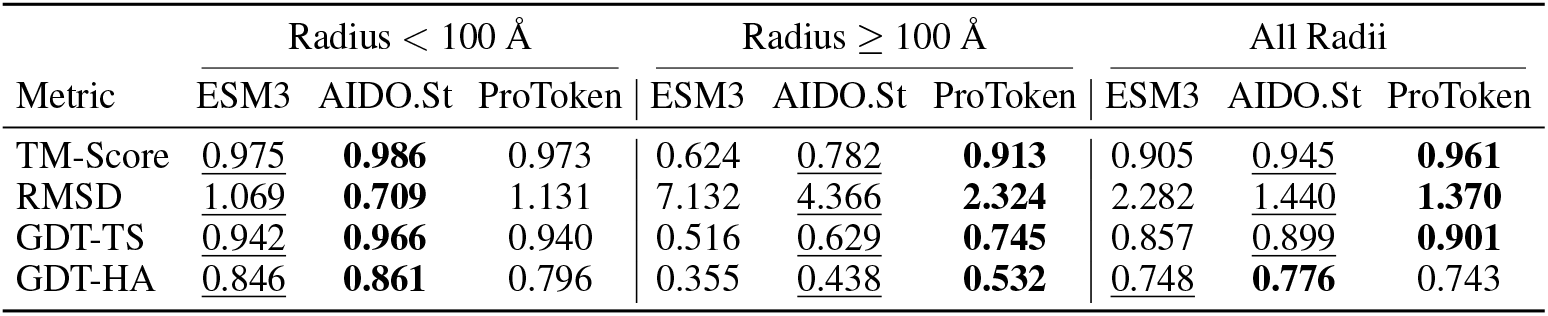
Comparison of structure reconstruction performance across different tokenizers for CASP15 protein structures, evaluated on proteins with radii below 100 Å and above 100 Å. Metrics include TM-Score(↑), RMSD(↓), GDT-TS(↑), and GDT-HA(↑), which measure the quality of reconstructed protein structures to their original forms. The results demonstrate that the overall performance of all tokenizers is quite good (TM-Score *>* 0.9), though large protein reconstruction remains quite challenging.

| Metric | Radius < 100 Å |  |  | Radius ≥ 100 Å |  |  | All Radii |  |  |
| --- | --- | --- | --- | --- | --- | --- | --- | --- | --- |
|  | ESM3 | AIDO.St | ProToken | ESM3 | AIDO.St | ProToken | ESM3 | AIDO.St | ProToken |
| TM-Score | 0.975 | **0.986** | 0.973 | 0.624 | 0.782 | **0.913** | 0.905 | 0.945 | **0.961** |
| RMSD | <u>1.069</u> | **0.709** | 1.131 | 7.132 | <u>4.366</u> | **2.324** | 2.282 | <u>1.440</u> | **1.370** |
| GDT-TS | <u>0.942</u> | **0.966** | 0.940 | 0.516 | <u>0.629</u> | **0.745** | 0.857 | <u>0.899</u> | **0.901** |
| GDT-HA | <u>0.846</u> | **0.861** | 0.796 | 0.355 | <u>0.438</u> | **0.532** | <u>0.748</u> | **0.776** | 0.743 |

### 4.2 Homology Detection Evaluation

In this task, we evaluate the effectiveness of the tokenization method by assessing its ability to capture structural similarities between proteins through homology detection. Specifically, we follow the same procedure as Foldseek [7], applying a Smith-Waterman alignment on tokenized sequences using the SCOPe40 database as shown in Fig.1.B. The performance shown in Table 4.2 is measured across three levels of structural homology: family, superfamily, and fold, using sensitivity up to the first error. This evaluation is crucial, as it allows us to test whether the tokenized representations retain essential biological relationships between proteins, despite their conversion into sequential discrete tokens. By maintaining the sequential information through tokenization, we ensure compatibility with protein language models (pLM) for future integration.

**Table 2:** Homology detection performance on the SCOPe40 database. Sensitivity up to the first error is measured at the family, superfamily, and fold levels.

|  | ESM3 | AIDO.St | ProToken | Foldseek |
| --- | --- | --- | --- | --- |
| (SCOPe40 retrieval) |  |  |  |  |
| Family | 0.741 | **0.795** | 0.703 | <u>0.769</u> |
| Superfamily | 0.343 | <u>0.413</u> | 0.329 | **0.438** |
| Fold | 0.026 | <u>0.056</u> | 0.052 | **0.072** |

Despite good reconstruction capacity, we realize that the highly global tokens of ProToken perform poorly to the highly local tokens of Foldseek (which, conversely, are unable to reconstruct structure). The iterations in AIDO.St were made to find a good compromise between the two.

### 4.3 Structure Prediction

To evaluate the effectiveness of our tokenization approach for multimodal training, we fine-tuned a protein-specific MoE language model with 16B parameters, AIDO.Protein [14], to predict structure-related tokens. These tokens are then input into the VQ-VAE decoder to reconstruct the full protein structure. This process allows us to assess how well our tokenized representations capture essential structural information and contribute to accurate structure prediction in downstream tasks. The alignment of the pLM with the token space is a key indicator of tokenization quality. The procedure involves tokenizing the PDB database and fine-tuning the language model to predict structural tokens based on amino acid sequences. This evaluation serves as the most robust method for assessing the quality of the tokens, as it directly measures their ability to align with amino acid-based models and support accurate structural reconstructions.

Due to the computational cost of fine-tuning models on large databases, we initiated this process fine-tuning ESM2-650M [13]. For this step, we utilized tokens generated from ESM3, ProToken, and our proposed model (see Table 3). Despite ProToken’s strong performance in reconstruction, it was less effective at predicting accurate structure tokens when integrated into a language model, as indicated by its lower TM-Score across all test sets. This highlights the limitation of focusing solely on reconstruction metrics, which can yield good structural representations but fail to generate tokens that are well-aligned to be used in a protein language model (pLM).

**Table 3:**
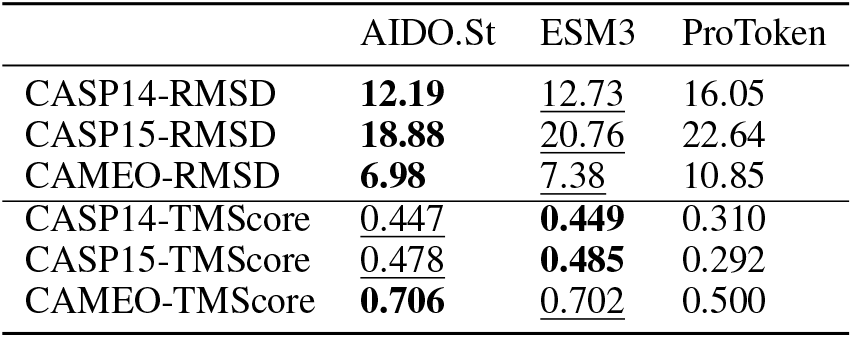
Comparing tokenizer quality through structure prediction performance. In these scenarios, we used ESM2-650M fine-tuned pLM for structure prediction.

|  | AIDO.St | ESM3 | ProToken |
| --- | --- | --- | --- |
| CASP14-RMSD | **12.19** | <u>12.73</u> | 16.05 |
| CASP15-RMSD | **18.88** | <u>20.76</u> | 22.64 |
| CAMEO-RMSD | **6.98** | <u>7.38</u> | 10.85 |
| CASP14-TMScore | <u>0.447</u> | **0.449** | 0.310 |
| CASP15-TMScore | <u>0.478</u> | **0.485** | 0.292 |
| CAMEO-TMScore | **0.706** | <u>0.702</u> | 0.500 |

In the second phase, we performed full fine-tuning of AIDO.Protein [14] to further improve structure prediction accuracy using our model. This more resource-intensive training allowed us to surpass the performance of the publicly released ESM3 model on all test sets, demonstrating the superior quality of our tokenizer (see Table 4). These results highlight the importance of high-quality tokenization in improving the overall performance of protein structure prediction models.

**Table 4:**
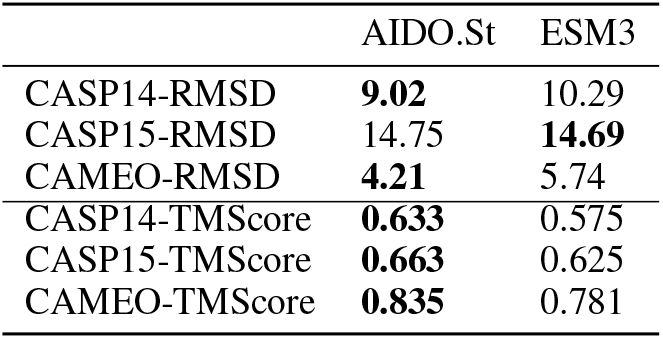
Final structure prediction performance comparison between our model and ESM3. We used the of-ficial ESM3 language model and tokenizer. Our model consists of a fully fine-tuned pLM based on our tokenization of the AFDB.

**Table 5:** Model hyperparameters for VQ-VAE.

| Component | Hyperparameters |
| --- | --- |
| Encoder | dim=128, layer=12, dropout=0.1 |
| Codebook | dim=384, vocab=512 |
| FoldingTrunk | dim_s=768, dim_z=128, layer=32, dropout=0.1 |
| Structure Module | dim_s=384, dim_z=128, layer=8, dropout=0.1 |

**Table 6:** Training hyperparameters for alignment experiment.

|  | ESM2 fine-tune | MoE fine-tune |
| --- | --- | --- |
| Training steps | 100k | 200k |
| Sequence length | 1024 | 1024 |
| Batch size | 128 | 2048 |
| Learning rate scheduler | linear decay | cosine decay |
| Warm up ratio | 0.01 | 0.025 |
| Max learning rate | 1e-4 | 1e-4 |
| Optimizer | Adam | Adam |

In a second step we did the costly full fine-tuning of our model in order to push further the structure prediction performance. This allowed us to reach higher performance than ESM3 released model on structure prediction. We outperformed ESM3 on all test sets, showing again the importance of the token quality and the higher performance of our tokenizer.

## 5 Discussion

In this work, we introduced AIDO.St, a tokenizer for protein structures, designed to capture critical structural information while preserving the sequential relationships with amino acids. This alignment is essential for multimodal tasks, such as integrating protein structure with sequence-based models. Visualization of the codebook space’s relation to secondary structure and amino acids usage reveals clear patterns in the codebook features (see Appendix C.3), suggesting potential improvements in the architecture. Evaluating tokenization quality is complex, as it requires balancing structural fidelity with downstream interpretability. Our assessments focused on two key tasks: reconstruction and homology detection. While reconstruction demonstrated the tokenizer’s ability to accurately represent protein structures, it alone is not sufficient. Good reconstruction may still result in tokens that are challenging for a language model to align with. Therefore, more robust assessments are necessary. We introduced homology detection to complement reconstruction by ensuring that the tokenization retains biologically relevant relationships—crucial for tasks involving multimodal integration.

Additionally, we explicitly trained a language model to predict these structural tokens for structure prediction tasks, and our model outperformed the publicly released ESM3. This further highlights the effectiveness of our approach in generating high-quality tokens that are both structurally faithful and interpretable by language models, advancing the potential for improved protein modeling.

## A Appendix: Training

### A.1 Training Data

For training the VQ-VAE, we utilized the PDB database released prior to May 1, 2022. The resolution threshold was set to 4Å, resulting in approximately 470k single chain structures. To reduce phylogenetic bias and ensure better sampling of the structural space, we applied sequence clustering with 40% sequence identity using MMseqs2^3^. This clustering was used to reweight the dataset during sampling, ensuring more balanced representation of diverse structures.

The data settings differed for the two language models. For the ESM2 model, we applied the same filtering settings as above but used a different time cutoff (May 1, 2020) to prevent data overlap with the CASP14 dataset. In contrast, the MoE model training leveraged both PDB and the AlphaFold Database (AFDB) to reduce overfitting. Proteins with an average plDDT score above 0.7 were selected from AFDB, resulting in 170M entries.

#### Data Sampling

During the training of the VQ-VAE, we addressed the underrepresentation of large proteins by oversampling structures with a large radius. The radius was defined as the maximum distance from the center of mass of all alpha carbons. We averaged the radius of each cluster obtained from MMseqs2 [15], and assigned the cluster-level sampling weight as min(max(10, radius), 90) − 10. During sampling, candidate clusters are first sampled according to the normalized sampling weights, and then one member is uniformly sampled from these clusters.

The ESM2 model was trained using uniform sampling from the PDB dataset. In contrast, the MoE model was trained with a balanced sampling strategy, where data was first sampled from the AFDB and PDB datasets in a 1:1 ratio, followed by uniform sampling within each dataset.

#### Data Cropping

During training of the VQ-VAE, the input structures are cropped to 256 residues following AlphaFold2’s training strategy, but we used a different cropping algorithm. The cropping process occurs as follows: Up to 10 random residues are selected as the initial candidates, and then neighboring residues are added iteratively based on their distance to the candidates. The distance threshold for adding neighbors is set to 15 angstroms. The process continues for a maximum of 10 iterations until the desired crop size is reached. If the structure contains fewer residues than the crop size, the entire structure is kept.

For the ESM2 and MoE models, we cropped the first 1024 tokens (including special tokens) to train the models. This approach simplified data processing and reduced computational costs; however, it introduced notable drawbacks. Cropping based on sequence order rather than structural context could exclude long-range structural dependencies, potentially limiting the models’ ability to capture global features. Additionally, this method disproportionately oversamples the prefix sequence, which may bias the models toward the beginning of the protein sequence. To address these limitations, we plan to explore cropping strategies that incorporate structural information and better balance sequence representation in future work.

### A.2 Architecture details

#### Structure Encoder

The encoder projects the backbone structure into a latent space where each residue is represented as a vector. The encoder is based on the Equiformer architecture [10, 16]. One good property of this model is that it leverages SE(3)/E(3)-equivariant features, enabling it to maintain rotational and translational symmetry. The to ensure that the output is independent of the input orientation, we discard the vector feature output at the final block.

The original Equiformer implementation is too memory demanding to run on proteins. Inspired by [16], we only used the first two orders of the spherical harmonics, and therefore, the hidden activations inside the encoder consist of scalar features and 3D vector features, which are invariant and equivariant to the input orientation, respectively. To ensure computational efficiency and maintain locality in the extracted features, the attention scope is restricted to the k-nearest neighbors in the input structure.

#### VQ-VAE Codebook

The latent vectors produced by the encoder are then quantized by mapping each vector to the nearest entry in a fixed-size codebook. This process converts the continuous latent vectors into discrete tokens, which are the same length as the amino acid sequence.

We apply a linear layer before the codebook and keep the feature dimension relatively small as suggested in [17] to encourage code utilization. Another trick for increasing code utilization is to random restart the unused codes. At the end of the training, there is no unused code.

During our experiments, we found that the codebook size (vocabulary size) is a critical hyperparameter that influences the granularity of the tokenization. Typically, a larger codebook size results in a better reconstruction performance. Since ProToken uses 512 codes, we also used 512 codes to make a fair comparison. It worths noting that ESM3 uses 8192 codes.

#### Structure Decoder

The quantized vectors are fed into a Multi-Layer Perceptron (MLP), which generates the “single embedding” input for decoder. Our decoder reuses the ESMFold Folding Trunk architecture. The Folding Trunk then reconstructs the full 3D structure, including side chains, based on the structural tokens.

The architecture is the same as used in ESMFold, but the number of layers and the hidden dimension are tuned. For simplicity, we disabled the recycling loop to save training and inference time. This architecture is very memory demanding, and we used techniques like gradient accumulation to avoid OOM.

#### Featurization

The backbone of a residue, composed of the N, CA, C, and O atoms, is represented using a rotation-translation approach, inspired by AlphaFold2. Since the backbone is generally rigid, its average coordinates **X**_ref_ ∈ ℝ^3×4^ can be computed from the dataset. During featurization, the input residue backbone **X** ∈ ℝ^3×4^ is aligned to **X**_ref_ using Kabsch alignment, resulting in **X** ≈ **RX**_ref_ + **t**. This equation means that knowing the rotation matrix **R** and translation vector **t** is sufficient to determine the original coordinates. For the orientation of **X**_ref_, we used N → CA as the direction of the *x* axis, and projected the direction of CA → C as the *y* axis using the Gram-Schmidt process. Besides rotation and translation, the residue’s offset (residue index) is also included in the features.

#### Embedding

Node embeddings are created by an Embedding layer for each residue. Since the model is agnostic to amino acid types, the Embedding layer has only one effective token, namely [MASK]. As required by the Equiformer architecture, the features are composed of scalar and vector components. A separate Embedding layer is used for the vector features, which also contains only one effective token. To ensure equivariance, the vector features are rotated by each residue’s rotation matrix.

For edge embeddings, the scalar part is derived from relative offset differences. These differences are clamped at 32, then transformed using a series of sinusoidal bases before being passed through an MLP layer. The vector part is based on the relative translation differences.

### A.3 Training Losses

For the reconstruction loss, we borrowed the losses from AlphaFold2 and used the OpenFold implementation. The borrowed losses are FAPE, supervised chi, and distogram loss following [2]. Besides these losses, we also added a translation loss and a distance map loss to stabilize the training.

For training the codebook, we followed the standard VQ-VAE training strategy. The VQ-VAE was trained with the straight-through estimator trick, and there is a commitment loss that encourages the encoder outputs to be close to the codebook vectors.

To encourage the robustness of the encoder output and the homology detection performance, we added a contrastive cropping loss, which encourages the latent similarity between 2 cropping.

#### Frame Aligned Point Error (FAPE) Loss

The FAPE loss is calculated by comparing the positions of atoms in the predicted structure (**x**_*j*_) with their corresponding positions in the true structure 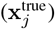, transformed into a common local frame (*T*_*i*_ and 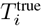). Each local frame is defined as the backbone frame of each residue.

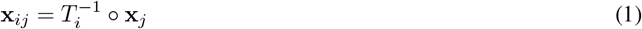

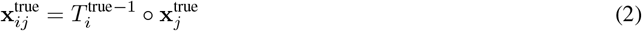

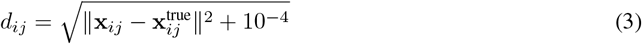

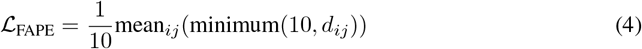

#### Commitment Loss

The commitment loss in a VQ-VAE encourages the encoder outputs to stay close to the embedding vectors from the codebook. It is defined as:

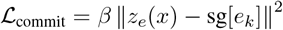

Where: *z*_*e*_(*x*) is the output of the encoder network for input *x*. *e*_*k*_ is the embedding vector from the codebook closest to *z*_*e*_(*x*). sg[·] denotes the stop-gradient operator, which prevents gradients from flowing through its argument during backpropagation. *β* is a hyperparameter that balances the strength of the commitment loss.

#### Translation Loss

FAPE optimizes structure similarity and is clamped by a 20Å threshold. In practice, we found the training a bit unstable. Here, we add another loss that optimizes the RMSD of the translation vectors.

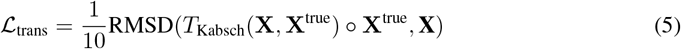

Here, *T*_Kabsch_(**X, X**^true^) is the Kabsch alignment from **X**^true^ to **X**, which are the translation vectors of all predicted and ground truth backbone frames. The Kabsch algorithm requires solving an SVD problem. To avoid singularity, we add a very small Gaussian noise to **X**.

#### Distance Map Loss

To further enhance training stability, we introduce the distance map loss, which enforces consistency in the pairwise distances between predicted and ground truth translation vectors. Let **x**_*i*_ and **x**^true^ represent the predicted and ground truth translation vector, respectively. We compare the pairwise Euclidean distances between all translation vector pairs for both the predicted and ground truth translations:

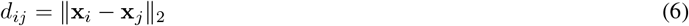

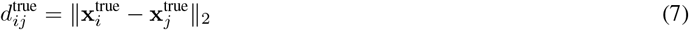

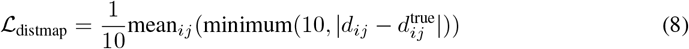

#### Cropping Loss

The cropping loss is a contrastive loss designed to encourage the preservation of latent embeddings when the input data is cropped. This loss ensures that even when parts of the input are removed or altered, the encoder still produces similar output.

Let **z**_*i*_ and 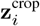 represent the per-residue quantized latent embedding of the original input and its cropped version, respectively. The cropping is done with the same algorithm mentioned in the “Data Cropping” paragraph. The distribution of the *i*-th code is defined via 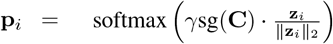 and 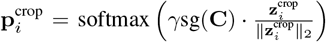, where sg stands for the stop-gradient operation, *γ* is a learnable temperature parameter that controls the sharpness of the distribution, and **C** represents the matrix of all codebook vectors. The loss is defined as the cross entropy between two distribution. Importantly, the loss only accounts for the common residues between the two crop views to ensure that only shared information is compared. The final cropping loss is given by:

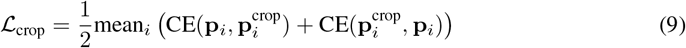

### A.4 Language Model Finetuning for Structure Prediction

We fine-tuned the pretrained protein language model to get the alignment model. Specifically, we denotes B as batch size, L as sequence length and D as latent dimension. Protein sequences (B, L) are input to language model to obtain the latent embedding (B, L, D). An additional two-layer MLP classification head (dimension D to 2560 to 512) is followed and predicts the precise structural token for every individual residual. The alignment model learns to project a sequence to corresponding structural token and cross entropy loss is employed.

During inference, protein sequence is converted to structural token with alignment model and fed into VQ-VAE decoder, together constructing a sequence-only protein structure predictor.

#### A.5 Model and training hyperparameters

The VQ-VAE architecture is set with the following hyperparameters, resulting in a 6M encoder and a 300M decoder.

The VQ-VAE was trained using the following hyperparameters: a cropping size of 256, batch size of 72, gradient clipping set to 0.1, Adam optimizer, learning rate warmup over 1k steps, a maximum learning rate of 1e-3, linear learning rate decay, a final learning rate of 5e-5 at 100k steps, weight decay of 1e-8, and actual training steps of 92k steps.

The alignment to the language model is trained with the following hyperparameters:

#### A.6 Training Curves and Convergence of the MoE Model

We tracked the training performance of the MoE model using TM-Score metrics on three datasets: CASP14, CASP15, and CAMEO. Figure 2 illustrates the training curves for these datasets. The curves were exported directly from Weights & Biases (wandb). The model demonstrates consistent improvement across all benchmarks; however, the steadily increasing trends indicate that the model has not yet converged. This suggests that further training could enhance performance.

**Figure 2.**
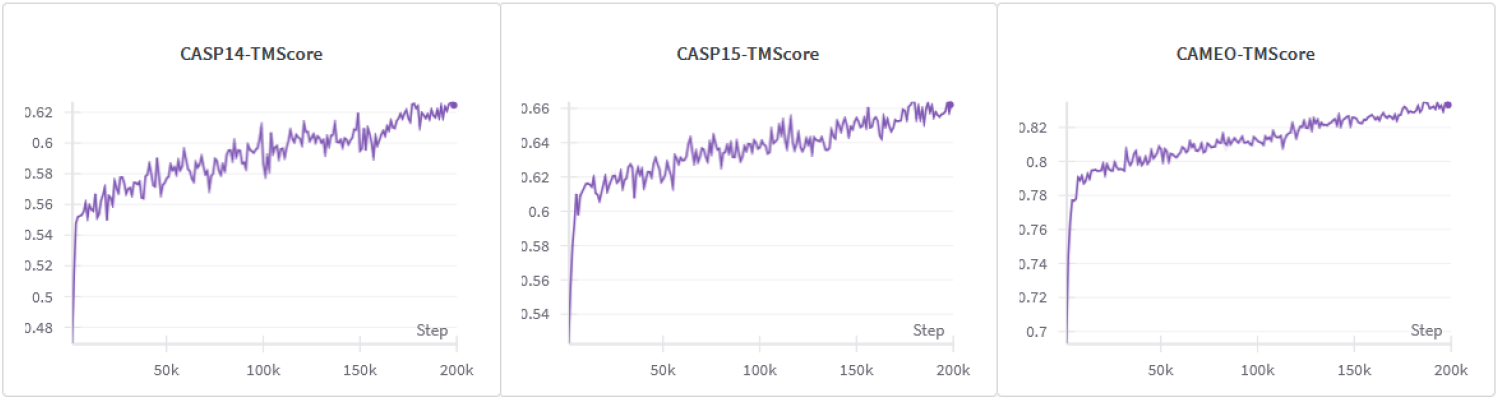
Training curves of the MoE model for CASP14, CASP15, and CAMEO datasets measured by TM-Score.

**Figure 3.**
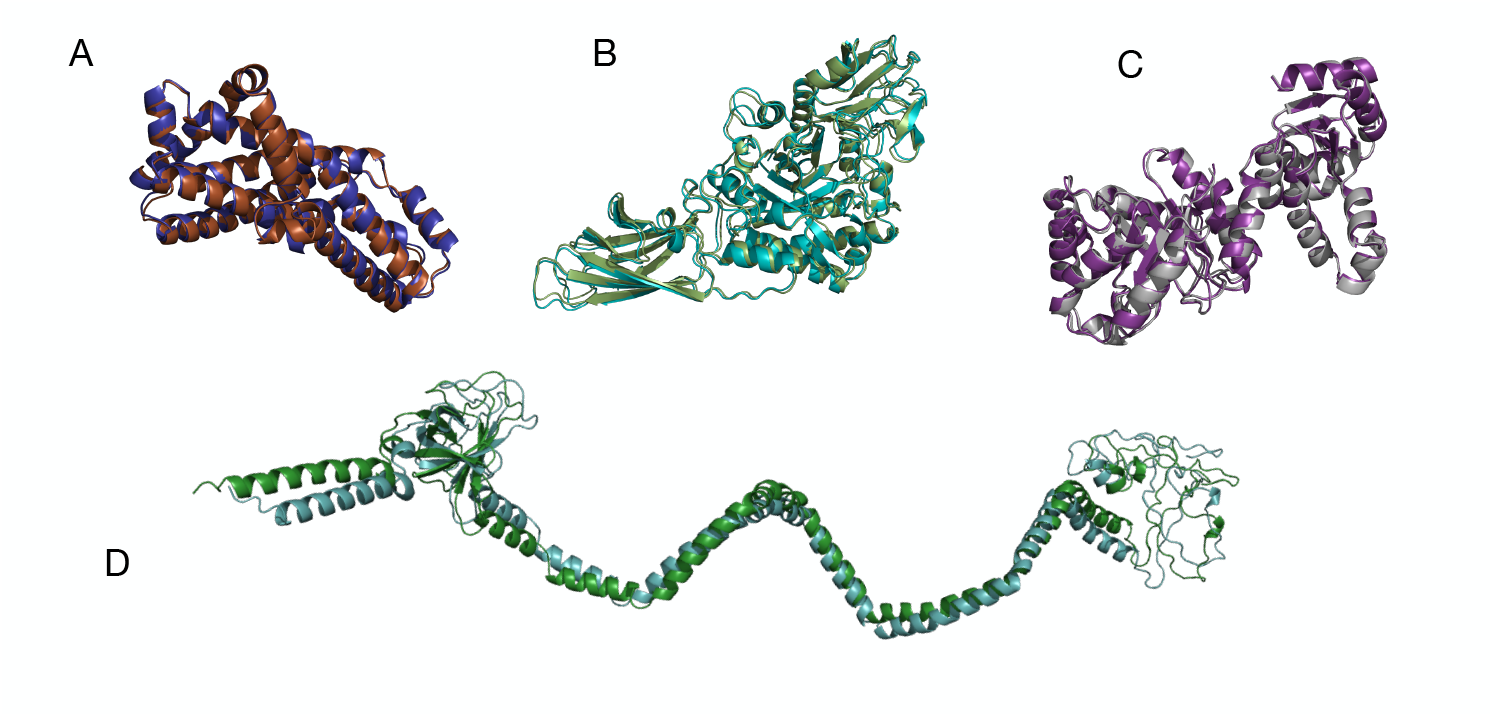
Reconstruction visualization of AIDO.St for different CASP15 entries. The entry D is an example of large radius proteins where the reconstruction performance are lower.A:T1137s9, B:T1188, C:T1180, D:T1137s6

**Figure 4.**
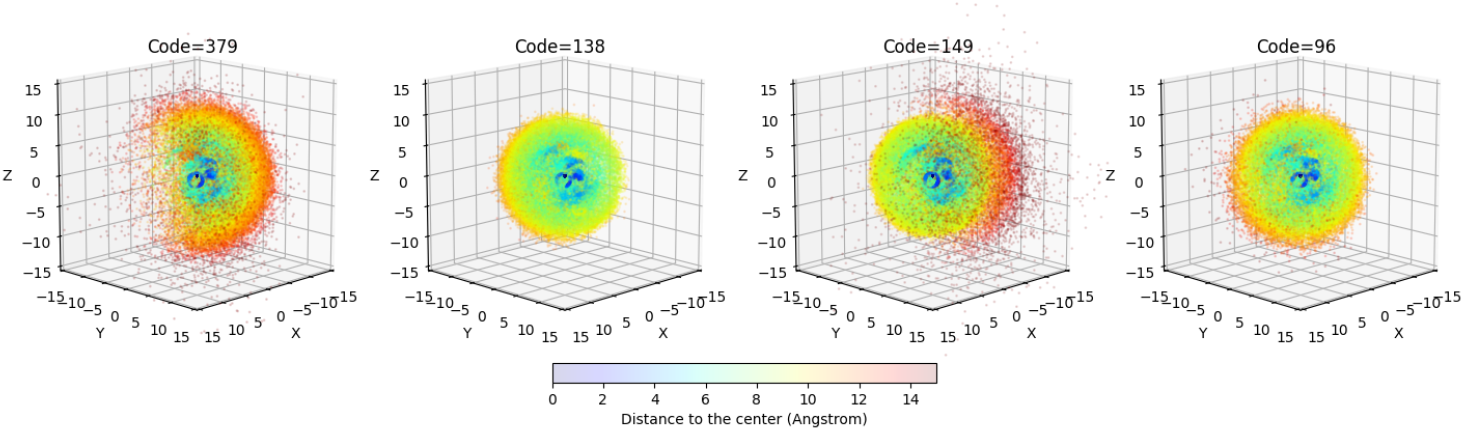
3D scatter plots visualizing the spatial distribution of neighboring residues for four different structure tokens. The plots show the relative positions of 30 nearest neighbors under the local frame, highlighting distinct density patterns for each structure token. The color of the points represents the distance from the central residue, with warmer colors indicating greater distances and cooler colors representing closer proximity. Points farther from the center exhibit greater distribution variety, reflecting the long-range interactions captured by the tokens. The distances are measured in angstroms.

**Figure 5.**
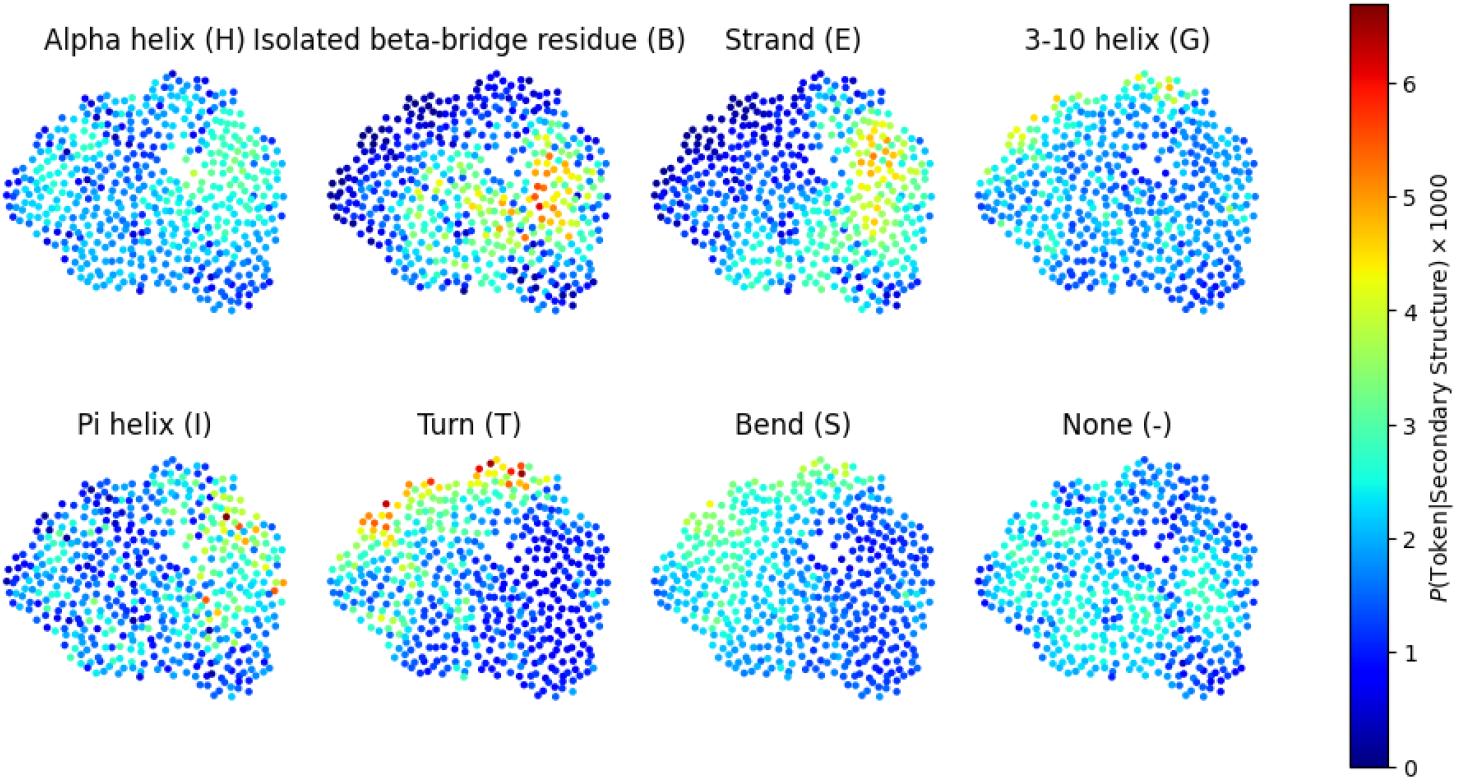
Visualization of tokens with respect to secondary structures. Each point represents a token from the codebook, mapped to different secondary structures, including Alpha helix (H), Isolated beta-bridge residue (B), Strand (E), 3-10 helix (G), Pi helix (I), Turn (T), Bend (S), and None (−). The location of each point is computed using t-SNE from the corresponding code vector. The secondary structures are computed using DSSP. The color scale represents the probability of each token given one specific secondary structure, with warmer colors indicating a higher probability. This visualization highlights the relationship between the learned token representations and various secondary structure types across the protein’s spatial configuration.

**Figure 6.**
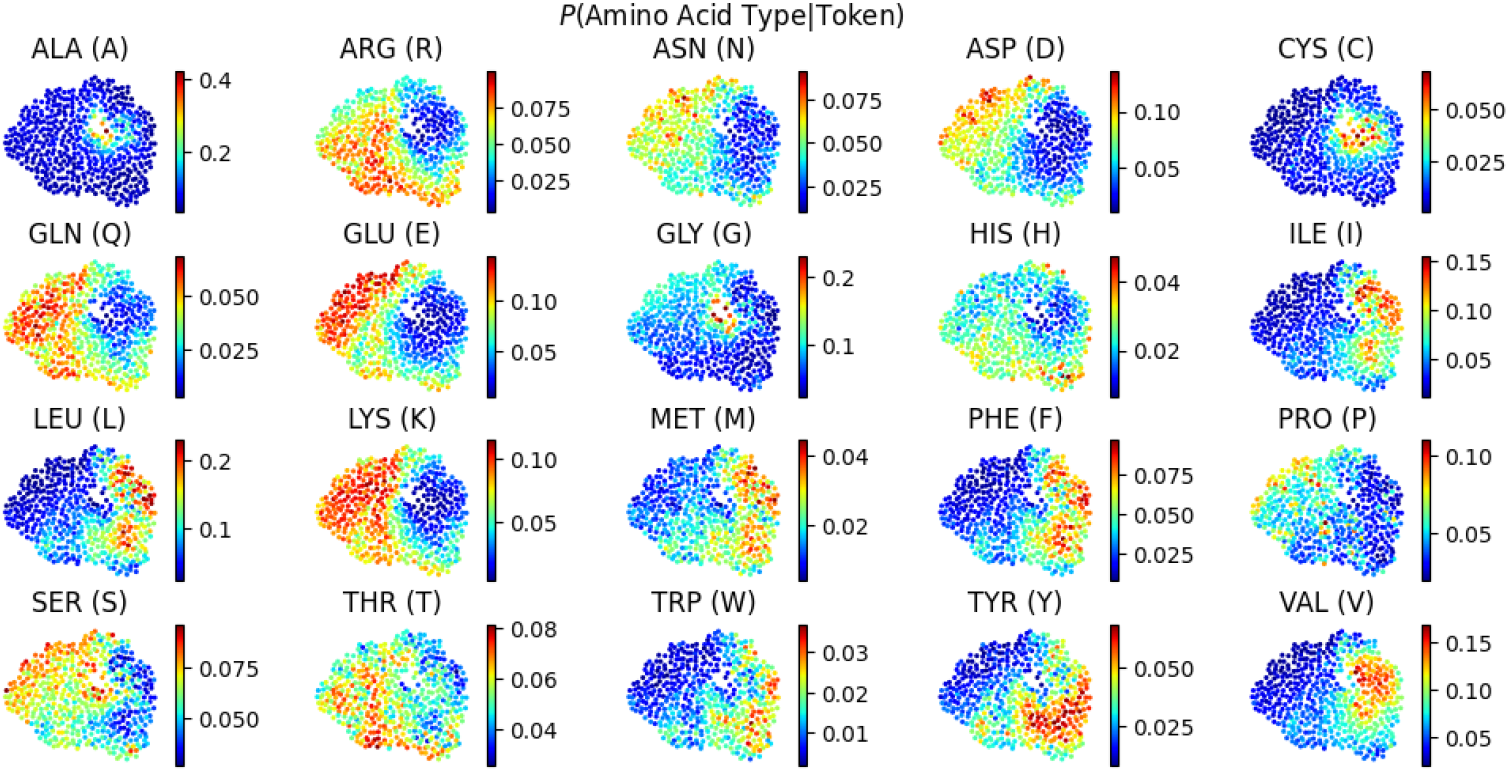
Visualization of tokens with respect to amino acid types. Each point represents a token from the codebook, mapped to different amino acid types. The location of each point is computed using t-SNE from the corresponding code vector. The color scale represents the probability of each amino acid type given one specific token, with warmer colors indicating a higher probability. This visualization highlights the relationship between the learned token representations and various amino acid types across the protein’s spatial configuration.

## B Appendix: Evaluation

### B.1 Reconstruction Metrics

#### TM-score (Template Modeling score)

TM-score is a measure of structural similarity between two protein structures. It evaluates how well the residues of one structure (typically a predicted model) match the residues of another structure (often the experimental structure) in 3D space. The TM-score ranges from 0 to 1, where a score closer to 1 indicates a higher similarity. Unlike RMSD, TM-score is not very sensitive to local errors and focuses more on the global topology.

The TM-score is calculated as:

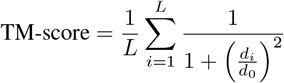

where *L* is the length of the protein, *d*_*i*_ is the distance between the i-th pair of residues, and *d*_0_ is a scale parameter (typically 1.24 Å times the cube root of *L* − 15).

Interpretation - A TM-score above 0.5 generally indicates a similar fold. - A TM-score below 0.2 indicates random similarity.

#### RMSD (Root-Mean-Square Deviation)

RMSD is a measure of the average distance between atoms (usually the backbone atoms) of superimposed proteins. It is sensitive to local structural differences, making it a good measure for comparing highly similar structures.

The RMSD is calculated as:

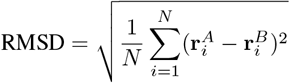

where *N* is the number of atom pairs, and 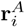 and 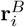 are the positions of the i-th atom in the two structures being compared. A superposition algorithm such as Kabsch that finds the best overlap between two structures is applied before computing RMSD. The best overlap is defined as the overlap that achieves minimum RMSD.

-Interpretation: - Lower RMSD values indicate higher structural similarity. - RMSD values are sensitive to the length of the protein and local differences, making them less ideal for comparing proteins of different sizes.

#### GDT-TS (Global Distance Test Total Score)

GDT-TS is a measure of the overall similarity between two protein structures, providing a percentage score that reflects the extent to which the structures are aligned. GDT-TS is less sensitive to local deviations than RMSD and offers a more holistic view of structural similarity.

-Calculation: GDT-TS is computed as the average of the percentages of residues that can be super-imposed within predefined distance cutoffs (1, 2, 4, and 8 Å). Specifically:

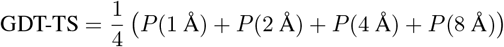

where *P* (*d* Å) is the percentage of residues with a distance less than *d* Å after optimal superimposition.

-Interpretation: - A higher GDT-TS indicates better structural similarity. - GDT-TS scores range from 0 to 100, with higher values representing a better match between the compared structures.

### B.2 Homology detection Metrics

The rationale is that a well-designed token representation will make proteins with similar structures and functions more alignable, resulting in higher alignment scores. We measure sensitivity up to the first false positive as our preferred metric. Alignments for each query are sorted by decreasing score and we count the number of hits above the first false positive. Sensitivity is then calculated as the total number of retained alignments divided by the number of related pairs in the database. When running Smith-Waterman, we used official substitution matrix for Foldseek, while we used cosine similarity as the substitution matrix for the rest of the models.

According to the setting of Foldseek, the retrieval on SCOPe40 is divided into 3 different subtasks, which are Family-level, Superfamily-level, and Fold-level. This division is based on the ontology of SCOPe dataset, where a tree structure categorizes proteins into Folds, Folds into Superfamilies, and Superfamilies into Families. For Family, Superfamily, and Fold subtask, false positives are defined as the proteins that are in a different Fold, and the true positives are defined as the proteins in the same Family, in the same Superfamily but not in the same Family, in the same Fold but not in the same Superfamily, respectively.

## C Appendix: Supplementary Results

### C.1 Examples of reconstruction

#### C.2 Examples of spatial distribution of neighboring residues

#### C.2 Visualization of tokens

We looked at interpretation of the codebook. To do so we started by ploting the t-SNE of the codebook embedding to better capture codebook similarity. Then we looked at the probability of these codebooks given some specific properties such as the secondary structure and the amino acid. Moreover we observe some interesting overlap, for example between 3-10 helix and Turn. Forcing the model to better separate codebook with different secondary structure, could introduce an interesting inductive bias in the training.

## D Data and Code Availability

We developed the ModelGenerator package to reproduce, apply, and extend the results in this manuscript https://github.com/genbio-ai/ModelGenerator.

Pre-trained models and finetuning data are also available on Huggingface at https://huggingface.co/genbio-ai.

## Footnotes

3 The clustering command is “mmseqs easy-cluster pdb.fasta pdb 40 tmp –min-seq-id 0.4 –threads 128”

